# A biochemical analysis of Black Soldier fly (*Hermetia illucens*) larval frass plant growth promoting activity

**DOI:** 10.1101/2023.01.06.523026

**Authors:** Terrence Green

## Abstract

Black Soldier fly (*Hermetia illucens*) larval (BSFL) frass separated from BSFL processed catering waste, and that recovered directly from larvae, was examined for its nitrogen, phosphate and potassium (N:P_2_0_5_:K_2_O), phytohormone and biogenic amine content, its plant growth promoting activity, and screened to test the hypothesis that bacteria characteristic of the genus *Enterococcus*, present in the biome of decaying catering waste and the larval gut, pass freely through the gut and are excreted in viable form into their frass. Its plant growth promoting activity was measured by comparing the growth of winter wheat berry (*Triticum aestivum*) grown in frass treated soil relative to that measured in untreated (control) soil. Its N:P_2_0_5_:K_2_O, biogenic amine and phytohormone composition were determined by standard soil analysis, HPLC and HPLC/GC-MS methodologies, respectively, and found to be too low to account for its plant growth promoting activity which induced a 11% increase in arial mass and shoot length in treated plants over controls. Colonies of *Enterococci* grew out on streaking frass collected directly from larvae on standard bile-esculin azide agar culture plates, confirming the hypothesis that viable *Enteroccoci* are excreted in their frass. Since *Enterococci* are capable of colonizing the rhizosphere and boosting the growth of plants on amendment into soil, these findings lend further insight into the underlying mechanism(s) accounting for the increased growth of plants growing in frass treated soils.

## Introduction

A number of researchers have conflated plant growth promoting activity associated with leftover waste byproducts recovered from Black Soldier fly (*Hermetia illucens*) larva (BSFL) processed feedstocks with BSFL frass even though it is generally well-known that much of the residual leftover byproduct, aside from frass, consists of leftover feedstock in varying stages of degradation, insect exuviate, and microorganisms [1–8]. Whereas there is no question that BSFL while feeding and growing off of feedstocks leave frass deposits in the feedstocks on which they feed, frass, separated free of contaminating residual leftover feedstocks processed by larvae has not heretofore been examined for its role in conferring to soil plant growth promoting activity upon its amendment into soil. Whereas it may be convenient to refer to the residual waste left behind by BSFL as frass, the residual waste’s biochemical and nutritional composition is not only complex but also varies depending on the source of the feedstock presented to larvae, how the feedstock waste was mixed and aerated as larvae processed the waste feedstock, the age and degree of decomposition of the waste feedstock at the time it was evaluated for its plant growth promoting activity on amendment into soil, etc. The complex makeup of BSFL processed waste recovered from BSFL bioreactors for all of these reasons presents a challenge to investigators interested in gaining a deeper insight as to the underlying mechanisms that may be set in motion on amending BSFL processed waste feedstocks into soil in conferring to the soil plant growth promoting activity.

This study was undertaken in an effort to gain more insight as to how and to what extent BSFL frass, itself, separated from and collected free of contaminating BSFL processed catering waste might be involved in conferring plant growth promoting activity to plants grown in frass treated soil, particularly, regarding the possible role of *Enterococci spp*. passing from the larval gut of BSFL into the frass which, upon amendment into soil, might plausibly account for the expression of plant growth promoting activity through colonization of the rhizosphere around the roots of receptive plants growing in frass treated soil. Plant growth promoting rhizobacteria capable of colonizing the roots of receptive plants, in particular, are present in decaying waste feedstocks [9–11]. While studying the soil biotic effects of residues of BSFL processed spent brewer’s grain feedstock amended into soil, Fuhrmann *et al*. [12] recently noted that there was a significant increase in microbial members of the genera *Bacillus* and *Streptomyces* in the biome of the BSFL processed spent brewer grain residues. The same two genera have also been identified in the gut biome of BSFL which include a number of strains having plant growth promoting rhizobacterial activity [13–15].

This study describes findings on examining the growth of winter wheat berry seedlings grown in soil treated with frass, its biogenic amine, phytohormone, and nitrogen-phosphate-potassium (N:P_2_0_5_:K_2_O) nutrient composition, and tests the hypothesis that *Enterococci* previously shown to be in relatively high abundance in the larval gut, are excreted and recoverable in viable form in frass collected directly from larvae washed free of contaminating waste feedstock. BSFL frass separated free of residual particulate BSFL processed catering waste on which BSFL have fed was found to contain only trace amounts of nutrients, biogenic amines and phytohormones, all at far too low of concentrations to account for its plant growth promoting activity observed upon amending the frass into soil. Numerous colonies of viable *Enterococci* were detected in frass collected directly from BSFL washed free of contaminating feedstock, confirming the hypothesis that *Enterococci* making up a significant portion of the larval gut are excreted into frass.

## Materials and methods

### Growth of BSFL, source of catering waste, recovery of catering waste leachate, and recovery of BSFL frass from BSFL processed catering waste and larvae

BSFL and frass separated from BSFL processed waste feedstock were obtained by growing BSFL in bioreactors loaded every fourth day with catering waste obtained from discarded cafeteria kitchen waste. Catering waste leachate was obtained by collection of the liquid fraction from unprocessed catering waste that drained freely from catering waste through a 20 mesh stainless steel screen.

BSFL frass was separated from BSFL processed catering waste in the same manner by collection of frass excreted by larvae that drained freely from BSFL processed catering waste through a 20 mesh stainless steel screen. Frass collected in this manner was analyzed for its dry matter nutrient N:P_2_0_5_:K_2_O content and compared to the nutrient N:P_2_0_5_:K_2_O content of leftover BSFL processed catering waste from which it was obtained.

BSFL frass collected directly from larvae was used in screening frass for the presence of viable *Enterococci spp*. on BEA agar plates and was obtained by removing larvae from catering waste on which they were feeding, washing catering waste completely free from their larval exoskeleton through repeated suspension and rinse cycles of the larvae in sterile water, and by then collecting their larval excrement after suspending the washed larvae in conical tubes tilted at an angle of approximately 20 degrees (hereafter referred to as “tilt tubes”).

A more detailed description on how the larvae and frass were obtained can be found in Supplemental Information, “S1 Collection of BSFL_ catering waste and frass.pdf”.

### Evaluation of BSFL frass plant growth promoting activity

Plant growth promoting activity was evaluated by measuring the effect of adding BSFL frass recovered from BSFL processed catering waste to potting soil placed in pots planted with winter wheat berry housed inside a greenhouse with diurnal exposure to natural sunlight and maintained at a constant temperature of 22 C. Control seedlings were set up in parallel by adding sterile water in place of frass to the potting soil. Frass (50 ml), was delivered to each of four pots containing 2.5 g seeds per pot started one week prior to the addition of frass to the potting soil, and once again in the following week. At the end of a 30-day growth interval, the upper plant masses of the experimental and control sets of plants were harvested at soil level, and the average upper plant masses ±1 standard deviation was calculated and compared to that of the corresponding control (untreated) set of plants. The average shoot length ±1 standard deviation for each test set was also calculated by measuring the shoot length of 137 shoots drawn at random from the frass-treated and control plants.

### Detection of *Enterococci* discharged into frass

BEA agar plates were streaked with wire loop samples drawn from the aqueous suspension of larvae immediately after the transfer of washed larvae into the tilt tubes (T=0, control sample, frass negative), and following subsequent recovery of pure frass excreted by larvae into in the aqueous suspension. After streaking out samples drawn from the tilt tubes in this sequence, plates were incubated at 35 C and examined for colony growth 24 and 48 hours later in screening for viable *Enterococcus spp*. colonies growing on the surface of the agar plate and concomitant generation of dark brown-black iron esculetin end product in the agar associated with the hydrolysis of esculin included in the agar medium [16]. BEA agar plates were obtained from Hardy Diagnostics.

### Analysis of frass biogenic amine content

Frass recovered in the liquid leachate draining free from BSFL processed catering waste, and leachate draining free from catering waste unprocessed by BSFL were each analyzed for biogenic amine composition by Trilogy Analytical Laboratory, Washington, MO by HPLC [17].

### Analysis of frass phytohormone content

Frass recovered in the liquid leachate draining free from BSFL processed catering waste, and leachate draining free from catering waste unprocessed by BSFL, were each analyzed for phytohormone content by Lifesible, Shirley, NY, by HPLC-MS analysis. Additional details on the analysis method and results can be found in Supplemental Information, “S2 Copy of phytohormone calculated data.pdf” and in “S3 HPLC GC-MS phytohormone analysis.xls”.

### Analysis of frass and BSFL processed catering waste N, K_2_O and available P_2_O_5_ content

Replicate samples of frass recovered in the liquid fraction draining free from BSFL processed catering waste, and BSFL processed catering waste left behind in the BSFL bioreactors, were each analyzed for N, K_2_O and available P_2_O_5_ by A&L Western Agricultural Laboratory, Modesto, CA by AOAC Official Method 993.13 (combustion method), Method 983.02 (updated to flame emission spectroscopy in place of flame photometry) and Method 960.03 (phosphorous available in fertilizers), respectively, from which the average N, K_2_O and available P_2_O_5_ content of each ±1 standard deviation was calculated.

### Statistical analysis of data sets

Statistical analysis of data sets in calculating the average of replicate samples, standard deviation (SD) and one-tailed t-tests were done using Microsoft Office 365 Excel, Version 2207 software.

## Results

Catering waste leachate recovered in the filtrate draining free through a 20 mesh stainless steel mesh screen constitutes the liquid fraction of decaying waste and was therefore analyzed for its biogenic amine and phytohormone content to establish a baseline level of these two classes of plant regulatory byproducts for comparative purposes relative to the corresponding levels subsequently measured in frass recovered from BSFL processed catering waste. Trace baseline levels of cadaverine, histamine, putrescine and tyramine were detected in catering waste leachate whereas phenylethylamine, spermidine, spermine and tryptamine, if present, were below the lower limits of detection and deemed insignificant (**Table 1**). Trace albeit insignificant amounts of indoleacetic, absicic and jasmonic acid, and methyl jasmonate, and gibberellin A1, A3, A4, and A7 phytohormones were likewise detected in catering waste filtrate yet nearly completely cleared in the frass as were all of the corresponding biogenic amines (**Table 1)**.

**Table 1.** Comparison of biogenic amine and phytohormone content of catering waste leachate versus BSFL frass recovered from larval-processed catering waste.

| ***Biogenic amines (ng/ml)*** | ***Catering Waste Leachate*** | ***BSFL Frass*** |
| --- | --- | --- |
| cadaverine | 7.1 | <5<sup>a</sup> |
| histamine | 13.7 | <5<sup>a</sup> |
| putrescine | 86.8 | <5<sup>a</sup> |
| tyramine | 9.5 | <5<sup>a</sup> |
| phenylethylamine, spermadine, spermine, tryptamine | <5<sup>a</sup> | <5<sup>a</sup> |
| ***Phytohormones (ng/ml)*** |  |  |
| indoleacetic acid | 50.4<sup>b</sup> | 0.36 |
| abscisic acid | 20.3<sup>b</sup> | 0.04 |
| jasmonic acid | 4.7<sup>b</sup> | 0.18 |
| methyl jasmonate | 0.14<sup>b</sup> | 0.02 |
| giberrellin A1 | 0.025<sup>b</sup> | 0.07 |
| giberrellin A3 | 0.014<sup>b</sup> | 0.01 |
| giberrellin A4 | 0.035<sup>b</sup> | 0.01 |
| giberrellin A7 | 0.014<sup>b</sup> | 0 |

<sup>a</sup> lower limit of detection <5 ng ml<sup>-1</sup>. <sup>b</sup> average of duplicate determinations.

The average dry matter N:P_2_0_5_:K_2_O content of BSFL processed catering waste recovered from BSFL bioreactors was 3.27 ± 0.21 % (±1SD, n=3), 3.14 ± 0.08 % (±1SD, n=3) and 4.45 ±-0.38 % (±1 SD, n=3), respectively. The calculated average N:P_2_0_5_:K_2_O oxide ratio rounded to the nearest tenth at 1.0:1.0:1.3 revealed furthermore that the BSFL processed catering waste residue left behind is relatively uniform in its nutrient content.

On the other hand, the average dry matter N:P_2_0_5_:K_2_O content of frass draining free from BSFL processed catering waste at 0.09 ± 0.07 (±1 SD, n=5), 0.04 ± 0.03 (±1 SD, n=5) and 0.32 ± 0.23 (±1 SD, n=5), respectively, was an order of magnitude less than that recovered in BSFL processed catering waste. Its calculated average N:P_2_0_5_:K_2_O oxide ratio (rounded to the nearest tenth) at 1.0:0.4:3.5, respectively, reveals a notable downward shift in its residual N and especially P_2_0_5_ content relative to its K_2_O content. Frass recovered from BSFL processed catering waste applied to potting soil nevertheless accelerated the growth of winter wheat berry seedlings planted in the potting soil. **Table 2** contrasts the growth of potted winter wheat berry seedlings grown in potting soil amended with frass to that of control seedlings grown in potting soil unexposed to frass. The results show that frass, though very low in nutrient (N:P_2_0_5_:K_2_O), biogenic amine and phytochrome content, enhances the growth of seedlings grown in soil treated with frass as evidenced by an average 11% increase in total upper plant mass relative to that observed for seedlings grown in control (untreated) soil concomitant with a statistically significant 11% increase in the average shoot length of the plants.

**Table 2.** Average 30 day upper plant mass and shoot length growth rate of Winter Wheat Berry seedlings grown in potting soil treated with BSFL frass *versus* untreated (control) potting soil.

|  | Average Upper Plant Mass<br>(g/pot) | Average Shoot Length<br>(cm) |
| --- | --- | --- |
| BSFL frass – treated soil test set | 13.9 ± 1.4 (±1 SD, n=4) | 31.1 ± 3.7 (±1 SD, n=137)<sup>a</sup> |
| Control – untreated soil test set | 12.5 ± 1.7 (±1 SD, n=4) | 28.1 ± 4.2 (±1SD, n = 137)<sup>a</sup> |
| Percent growth relative to control test set | +11% | +11% |
| One tailed t-test | (0.17) | (1.5 x 10<sup>-9</sup>)<sup>a</sup> |

<sup>a</sup> Statistically significant findings.

Numerous microbial colonies characteristic of the genus *Enterococcus* grew out on BEA agar petri dish plates streaked with larval frass samples recovered directly from larvae washed free of contaminating waste feedstock before collection of the frass screened for *Enterococci* (**Fig 1**).

**Fig 1.**
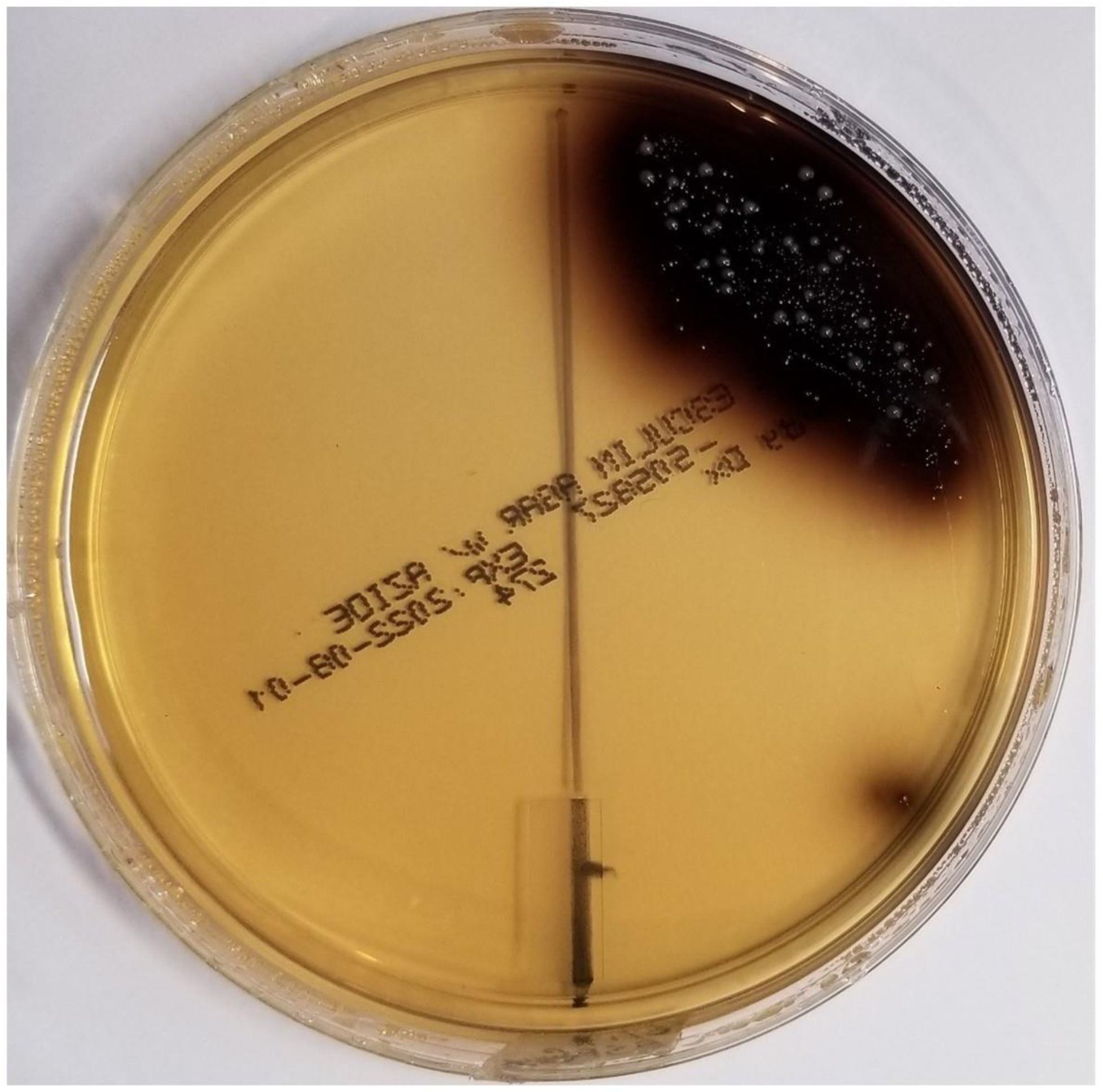
BEA agar plate assay confirming the presence of microorganisms characteristic of the genus *Enterococcus* in frass recovered from a tilt tube suspension of BSFL diluted 100-fold. Right side of plate – frass recovered after resuspension of washed larvae in water; Left side of plate - corresponding water control used in resuspending the larvae plated out on the same plate. Note detection of multiple colonies and positive iron esculetin coloration on right side of plate and the absence of colonies growing on the left side of the plate.

**Fig 2** shows the results of plating out a corresponding control sample drawn from a tilt tube before the larvae were able to excrete a significant amount of frass into the tilt tube (T=0). Only two positive colonies were detected in the control indicating that the preponderance of *Enterococci* detected as in **Fig. 1** were from frass left behind in the tilt tube following sufficient time after the transfer of larvae into the tilt tubes for the larvae to leave visible deposits of frass in the aqueous suspension.

**Fig 2.**
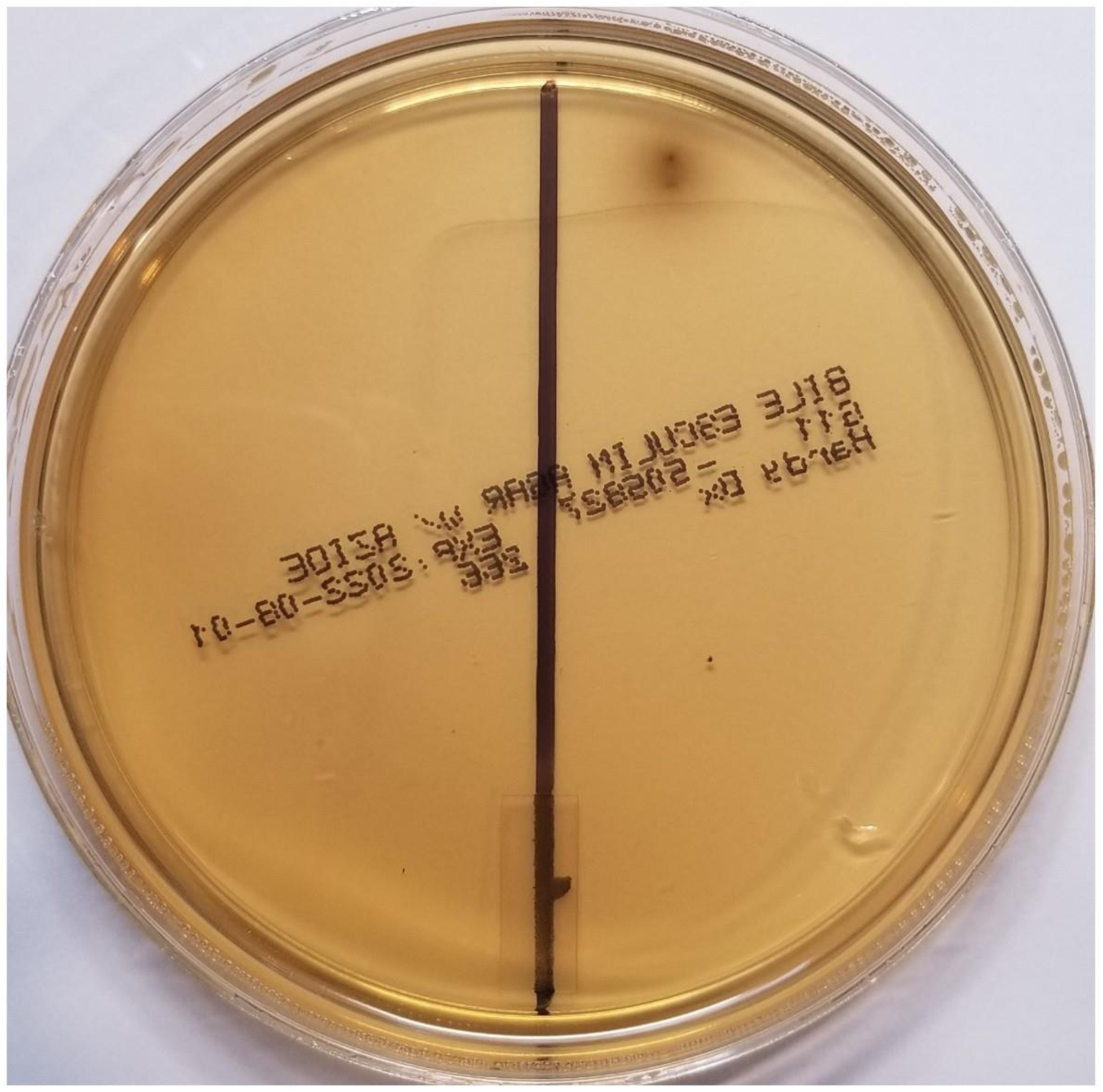
BEA agar plate assay for microorganisms characteristic of the genus *Enterococcus* drawn immediately after resuspension of washed BSFL in sterile water. Right side of plate – sample drawn from washed larvae immediately after resuspending the larvae in water (T=0); Left side of plate - the corresponding negative water control.

## Discussion

Gärttling and Schulz [1], on reviewing and collating recent published literature on the use of BSF frass as a commercial fertilizer, tacitly acknowledged in their review that “In the context of black soldier fly (BSF) rearing, often the residues from production – mainly faeces but also undigested substrate – are addressed as frass in a broader sense.” Klammsteiner et al. [13] also noted the same issue regarding the use of the term frass with regard to BSFL research on this subject in stating that “Frass in general describes insect excretions, but in a commercial context it often refers to a mixture of mainly insect feces, substrates residues and shed exoskeletons.” Lopes, Yong and Lalander [2], in yet another critical review, commented on the instability of the byproduct referenced in several of the publications they reviewed in noting published reports showing both positive and negative effects of the frass byproducts regarding its capacity to confer to soil plant growth promoting activity, differences in published C/N ratios and NPK content among the data sets published by different authors. Given the limitations in the definition of what actually constitutes BSFL frass regarding the quality and composition of source material on which BSFL greenhouse and field trial conclusions have been drawn, particularly concerning its biochemical and microbial attributes as a nutrient fertilizer, an examination of the properties of frass, itself, is overdue. Frass collected as in **Materials and Methods** was examined to determine: (i) its percent dry weight N:P_2_0_5_:K_2_O content which if significant could be serving as a source of nutrient fertilizer in promoting the growth of a plant; (ii) its phytohormone content, particularly its indole acetic acid and/or gibberellin content, which if present in sufficient quantities could account for a growth spurt in plants due to the action of phytohormones present in the frass inducing stem cell elongation and/or an overall increases in a plant’s areal mass [18–20]; (iii) its biogenic amine content, which if in significant quantities could confer to a plant an advantage in growth by alleviating abiotic stresses [21–24]; (iv) whether or not upon amendment into soil it confers to soil plant growth promoting activity; and (v) whether or not it contains *Enterococci spp*. in viable form which if present could account for its plant growth promoting activity. The percent dry matter N:P_2_0_5_:K_2_O and oxide ratio of BSFL processed catering waste without separating frass from the residual leftover waste byproducts reported in this study averages 3.3:3.1:4.5 and 1.0:1.0:1.3 (rounded to the nearest tenth), respectively, which is in reasonably good agreement with the data sets collated and the calculated average oxide ratio of 1.0:0.9:1.1 summarized by Gärttling and Schulz [1]. The percent dry matter average N:P_2_0_5_:K_2_O ratio of frass separated free of BSFL processed catering waste, on the other hand, at 0.09:0.04:0.32 (reported to the nearest hundredth), and very low phytohormone and biogenic amine content (**Table 1**), are all insignificant. These latter results make it very unlikely that the plant growth promoting activity reported here on amending frass into soil (**Table 2)** is attributable to its N:P_2_0_5_:K_2_O, biogenic amine and/or phytohormone content.

While a direct N:P_2_0_5_:K_2_O, biogenic amine or hormonal effect accounting for the expression of frass plant growth promoting activity seems implausible based upon the chemical composition of the frass, there is ample evidence that decomposing catering waste contains within its biome a number of plant growth promoting rhizobacteria which on colonizing the roots of susceptible plants could have a significant role in solubilizing and securing soil nutrients, fixing N into the soil, and in regulating plant growth through in situ production of plant regulatory hormones [9–11,25]. The presence of microorganisms in the phylum Firmicutes, including those from the genus *Enterococcus* as shown here in this study, and from the genus *Clostridium* also identified in food waste processed by BSFL (26), ingested by BSFL feeding off of the waste feedstock, passing through and colonizing the larval gut, and then excreted in viable form as frass, is of particular interest since there is evidence that several microorganisms from these two genera promote on amendment into soil the growth of barley and mung bean plants, and an enhancement in seed germination rates for beet, barley, wheat, red radish, cucumber and tomato plants [27,28] and Pakchoi (*Brassica rapa* L.) [29].

Sasmita et al. [30] and Pakpahan et al. [7] have also published data following treatment of coca seedlings and sugar cane, respectively, demonstrating a link between the expression of plant growth promoting activity and the presence of plant growth promoting activity in liquid BSFL extracts recovered from BSFL processed vegetal, fruit and dairy waste feedstocks which they referred to as a liquid biofertilizer. Microorganisms identified in the liquid biofertilizer capable of fixing N in soil, P solubilizers and phytohormone producers were associated with *Pseudomonas spp*., *Bacillus spp*., *Trichoderma spp*., *Streptomyces spp*., *Azotobacter spp*., and *Azospirillum spp*.

Since plant growth promoting rhizobacteria are clearly present and grow in decaying waste, it seems reasonable to expect that leftover BSFL processed waste feedstocks, amended back into soil, ought to similarly exhibit plant growth promoting activity on amendment back into soil as a direct result of the introduction of those plant growth promoting rhizobacteria already present in the leftover BSFL processed waste on amendment into soil colonizing the rhizosphere of receptive plants growing in treated soils.

It is possible that the plant growth promoting activity reported here on amendment of frass recovered from BSFL processed catering waste into soil (**Table 1**), and that referred to as a liquid biofertilizer in the studies published by Agustiyani et al. [29], Pakpahan et al. [7] and Sasmita et al. [30], could be due to carryover of plant growth promoting rhizobacteria also present in the leftover waste byproducts from which the liquid/frass fraction was derived. However, the BEA agar petri dish results of this study (**Figs 1** and **2**) confirm that *Enterococci spp*. are present in frass and that their presence is not due to cross contamination since in this instance the frass was collected directly from larvae washed free of contaminating waste feedstock before screening of the frass for *Enteroccoci* ensued.

This study does not imply that the plant growth promoting activity conferred to soil on amending frass into soil is due only to the presence of *Enterococci* colonizing the rhizosphere of plants growing in treated soils. Other rhizobacteria not yet identified may also be passing from the larval gut into the larval frass which could also on amendment of the frass in soil colonize the roots of receptive plants. Further studies must therefore still be done to fully map out how frass separated free of residual BSFL processed waste feedstocks, amended back into soil, affects the soil biome leading up to a boost in the growth of plants growing in treated soils.

In summary, confirmation of the hypothesis that rhizobacterial microbes characteristic of *Enterococci* previously identified as part of the larval gut biome are excreted into their frass provides a plausible explanation accounting for the plant growth promoting activity of frass observed on amending processed catering waste and/or frass into soil. Given the growing awareness and application of rhizobacteria as an effective means of boosting agricultural crop yields, further research aimed at identifying the relative abundance, diversity and characteristics of rhizobacteria passing from the larval gut biome into frass is warranted and ought to prove very helpful in gaining a better understanding as to how best to recycle this BSFL waste byproduct generated by the BSFL industry which in the long run could be used in agricultural applications to boost crop yields while possibly offsetting the use of chemical fertilizers applied to soils.

## Acknowledgements

The author is grateful to the administrative staff and cafeteria personnel at Lewis and Clark College for allowing the author free access to catering waste used in this study.

## Supporting Information

**S1 Collection of BSFL_BSFL processed catering waste and frass.pdf**

**S2 Copy of phytohormone LZBD12292005 Calculated data.pdf**

**S3 HPLC GC-MS phytohormone analysis.xls**

