## Supplementary material for "A biochemical analysis of Black Soldier fly (*Hermetia illucens*) larval frass plant growth promoting activity": S1 Collection of BSFL_ catering waste and frass (from another version of this preprint: version 1, 2023-01-07)

Author: Terrence R. Green

Supplemental Information, “S1 Collection of BSFL_ BSFL processed catering waste and frass.pdf”.

*Collection of BSFL and BSFL processed catering waste*.

BSFL were grown from egg clutches deposited by female BSF in catering waste mixed with shredded wood chip bulking agent under steady-state conditions in 400 x 600 mm plastic bioreactors fabricated out of plastic totes having 0.6 cm holes spaced approximately 10 cm apart on all four vertical walls of the totes to provide for adequate ventilation and aeration of the waste feedstock added to the bioreactors. Catering waste was loaded into the bioreactors every fourth day at a loading rate of 5 Kg catering waste mixed with approximately 0.07 Kg shredded wood chips per square meter bioreactor space per day.

The bioreactors were housed inside a 24 ft x 12 ft shed designed to allow natural daylight to pass through skylights installed in the roof of the shed by which adult BSF were able to freely mate and deposit their egg clutches in the catering waste. The ambient temperature of the shed was maintained between 25 and 30 C and kept at a relative humidity year-round between 55 to 75%.

BSFL growing off the catering waste were collected in “K” type rain gutters secured around the perimeters of the bioreactors on reaching the prepupa stage in their life cycle and then set aside in plastic trays to allow the collected prepupae to pupate and emerge from their puparia as adult flies. The adult flies mated and reseeded incoming catering waste with new egg clutches in sustaining the output of BSFL.

BSFL processed catering waste was obtained by drawing off approximately 10% of the catering waste accumulating in the bioreactors infested with young larvae approximately every 30 days and setting aside the waste in separate bioreactors which were kept for an additional approximate 30 days inside the shed over which time most larvae exited from the latter bioreactors. During this holding period no catering waste was loaded into the latter bioreactors.

*Collection of BSFL frass*.

Frass screened for *Enterococci* on BEA agar plates was collected directly from BSFL by removing larvae at about two-weeks of age from the catering waste on which they were feeding and washing them on stainless steel 20 mesh screens with copious quantities of tap water to rid the larvae of residual waste otherwise carried over on their exoskeleton, and by then resuspending the larvae in clean tap water in plastic conical tubes tilted at an angle of approximately 20 degrees covered with mosquito netting at a larval concentration of five larvae per ml. The tilt tubes were maintained at a constant temperature of 22 C throughout the period in which frass was collected. **Figure 1** shows the experimental setup used with the tilt tubes in collecting frass used in screening for *Enterococci*.

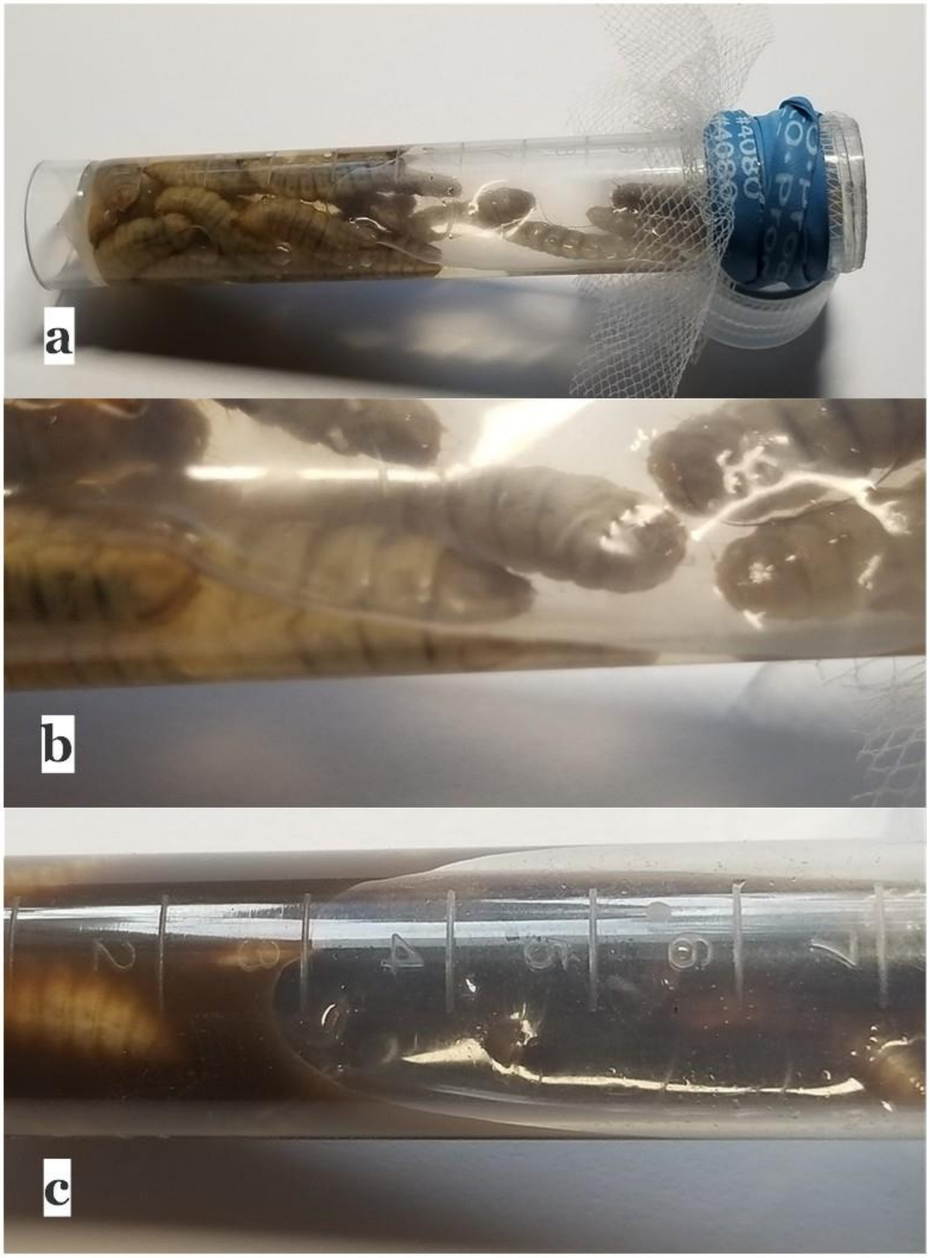


**Fig. 1**. Collection of BSF frass following the washing of larvae fed catering waste and resuspension of the washed larvae in conical tilt tubes in water. **Figure 1a**, the appearance of larvae suspended in water immediately after washing their larval exoskeleton free of food waste debris; **Figure 1b**, the appearance of the aqueous fraction

approximately 2 minutes after resuspension of the washed larvae in water; **Figure 1c**,the appearance of the aqueous fraction approximately 6 hours after suspension of the washed larvae revealing presence of significant amounts of frass excrement in the aqueous suspension. Photos in **Figures 1b** and **1c** are shown at approximately 4X magnification over that of **Figure 1a**.

Frass used in the plant growth promoting experiments was obtained from the BSFL processed catering waste by slurring the processed waste at a ratio of 1 Kg per 20 L of tap water and filtering the aqueous slurry through a 20-mesh stainless steel screen.
