## Supplementary material for "A biochemical analysis of Black Soldier fly (*Hermetia illucens*) larval frass plant growth promoting activity": S2 Copy of phytohormone calculated data (from another version of this preprint: version 1, 2023-01-07)

www.lifeasible.com
Lifeasible
17 Ramsey Rd. Suite 209
Shirley, NY 11967

### PROJECT REPORT

#### Plant Hormone Analysis for Biomass Fermented Waste

**Project Number LZBD12292005**

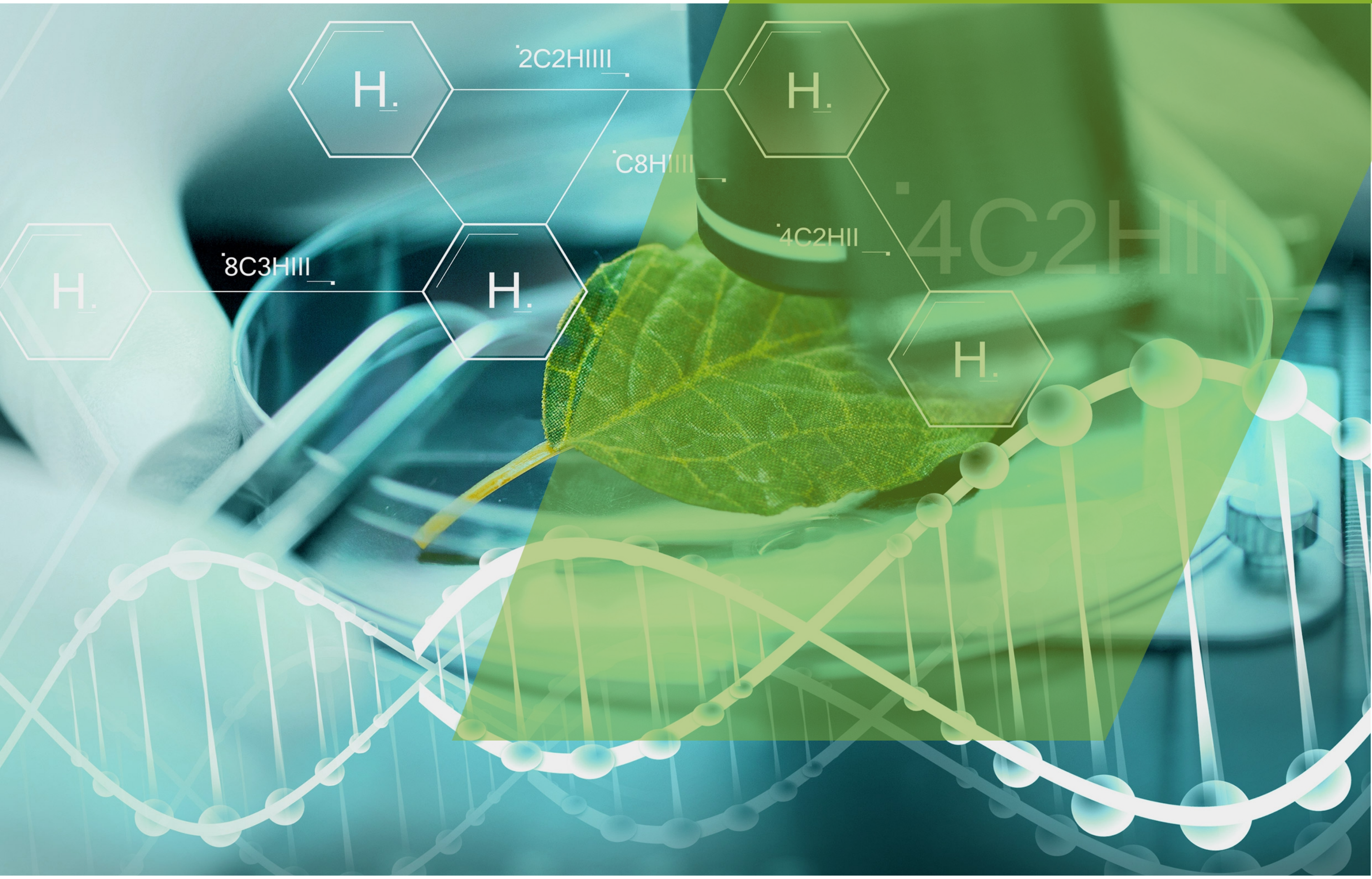

Report generated by
Rujane Zhang Ph.D., R&D Manager

Report Reviewed by
Miao Li Ph.D., Project Manager

On February 8 2021

### 1 Project Information

The service quotation [Number LZBD12292005] is for quantification analysis of the plant hormones for biomass fermented waste using electrospray ionization-high-performance liquid chromatography tandem mass spectrometry (ESI-HPLC-MS/MS). The plant hormones include indole-3-acetic acid (IAA), jasmonic acid (JA), methyl jasmonate (Me-JA), abscisic acid (ABA), gibberellic acid 1 (GA1), gibberellic acid 3 (GA3), gibberellic acid 4 (GA4), and gibberellic acid 7 (GA7). Two samples derived from biomass fermented waste are provided by customer. We have successfully finished the project.

### 2 Methods and Materials

#### 2.1 Samples

- Two samples of BSF101E and FS101C are provided by customer.

| Sample Label | Sample NO. | Form |
|---|---|---|
| BSF101E | B1 | Liquid |
| FS101C | B2 | Liquid |

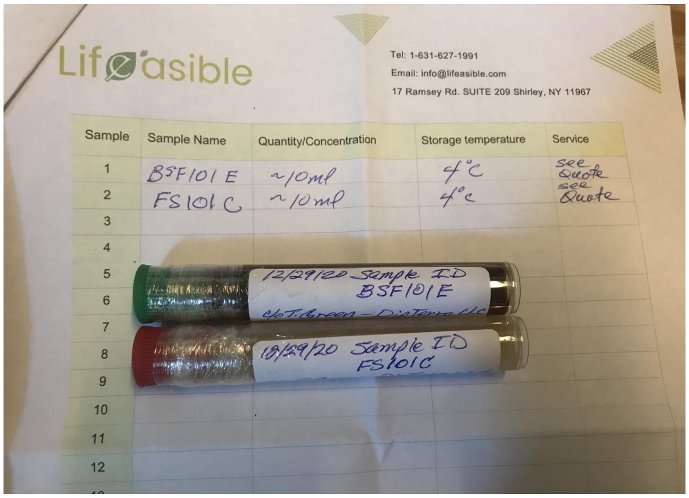

#### 2.2 Reagents and Consumables

- Indole acetic acid (IAA) standard (Sigma-aldrich)
- Abscisic acid (ABA) standard (Sigma-aldrich)
- Jasmonic acid (JA) standard (Sigma-aldrich)
- Methyl jasmonate (Me-JA) standard (Sigma-aldrich)
- Gibberellin acid 1 (GA1) standard (TRC)
- Gibberellin acid 3 (GA3) standard (Sigma-aldrich)
- Gibberellic acid 4 (GA4) standard (Sigma-aldrich)
- Gibberellic acid 7 (GA7) standard (TRC)
- Deuterium-ABA standard (Sigma-aldrich)
- Deuterium -IAA standard (Sigma-aldrich)
- Chromatographic methanol (Merck)

- Analytical acetonitrile (Merck)
- Chromatographic formic acid (Merck)
- C18 Queschers packing (Anpel)
- PSA Queschers packing (Anpel)

#### 2.3 Equipment

- AGLIENT1290 HPLC system (AGLIENT)
- AB SCIEX-6500 Qtrap MS/MS (Applied Biosystems)

#### 2.4 Methods

##### 2.4.1 Preparation of Standard Solution of Plant Hormones

1) 984 μL methanol was added into a 1.5 mL centrifuge tube and 500 μg/mL standard reserve solution of 2 μL for each hormone was added. The solution was well mixed prepared as the mother solution with final concentration of 1 μg/mL.
2) 996 μL methanol was added into a 1.5 mL centrifuge tube and 2 μL of 500 μg/mL internal standard reserve solution of each hormone was added. The solution was well mixed and prepared as the mother solution with final concentration of 1 μg/ mL.
3) Methanol of 989.9 μL, 989.8 μL, 989.5 μL, 988 μL, 985 μL, 970 μL, 940 μL, 790 μL was added into a 1.5 mL centrifuge tube, and the mother solution prepared in step (1) was added into the methanol solution in sequence of 0.1 μL, 0.2 μL, 0.5 μL, 2 μL, 5 μL, 20 μL, 50 μL, 200 μL, respectively. Then add 10 mL the internal standard in step (2) per tube to make standard curvature solution with final concentration 0.1 ng/ mL, 0.2 ng/mL, 0.5 ng/mL, 2 ng/mL, 5 ng/mL, 20 ng/mL, 50 ng/mL, 200 ng/mL and containing 10 ng/mL internal standard.

##### 2.4.2 Phytohormones Extraction

1) Shake the liquid sample thoroughly, accurately take 5 mL into the tube, and add 10 mL dichloromethane and 2 μL internal standard mother solution.
2) The sample solution was briefly vortexed and then kept at 4 °C overnight.
3) Following centrifugation of 5 min at 12,000 g at 4 °C, the lower dichloromethane phase was taken.
4) Add 35 mg C18 packing, shake violently for 30 s, centrifuge at 10,000 g for 5 min, and get the supernatant.
5) The supernatant was dried by nitrogen and then dissolved in 200 μL methanol.

6) After filtration through a membrane filter (0.22 μm), 2 μL extract was injected onto a C18 column mounted on an analytical HPLC system (AGLIENT) equipped with AB SCIEX-6500 Qtrap MS/MS.

#### 2.5 HPLC-MS/MS Program

##### 2.5.1 HPLC Conditions for Phytohormones

Column: Poroshell 120 SB-C18 reversed phase column (2.1 mm x 150 mm, 2.7 μm);

Column temperature: 30 °C;

Mobile phase: A:B = (Methanol / 0.1% formic acid): (water / 0.1% formic acid);

Gradient Parameters of HPLC:

**Table 1 Gradient Parameters of HPLC for Phytohormones**

| Time (min) | Flow Rate (mL/min) | A% |
|---|---|---|
| 0.1 | 0.3 | 20 |
| 1-3 | 0.3 | Increment to 50 |
| 3-9 | 0.3 | Increment to 80 |
| 9-10.5 | 0.3 | 80 |
| 10.5-10.6 | 0.3 | Decrease to 20 |
| 10.6-13.5 | 0.3 | 20 |

Injection volume: 2 μL.

##### 2.5.2 Mass Spectrometry Parameter for Phytohormones

Ionization mode: ESI positive and negative ion mode monitoring;

Scan type: MRM;

Air curtain gas: 15 psi;

Spray voltage: +4500 v, -4000 V;

Atomizing gas pressure: 65 psi;

Auxiliary gas pressure: 70 psi;

Atomization temperature: 400 °C.

**Table 2 Selected Reaction Monitoring Conditions for Protonated or Deprotonated Phytohormones ($[M+H]^+$or$[M-H]^-$)**

| Name | Polarity | Parent Ion (m/z) | Daughter Ion (m/z) | Declustering Voltage (V) | Collision Energy (V) |
|---|---|---|---|---|---|
| IAA | + | 176.2 | 129.8*/102.9 | 65 | 12/42 |
| JA | - | 209.2 | 58.9* | -54 | -16 |
| MeJA | + | 225.1 | 151.1/193.1/133* | 50 | 16/10/18 |
| ABA | - | 263.1 | 153.0*/204.2 | -60 | -14/-27 |
| GA1 | - | 347.2 | 259.2*/273.0 | -25 | -28/-16 |
| GA3 | - | 345.2 | 143.0/239.0* | -80 | -30/-33 |
| GA4 | - | 331.1 | 243.2*/213.1 | -131 | -24/-39 |
| GA7 | - | 329.2 | 223.2/241.1* | -89 | -38/-22 |

*Quantitative ions

### 3 Results

#### 3.1 Calibration Curves

Each calibration curve was drawn by AB SCIEX-6500 Qtrap MS/MS instrument according to data of STD1-8. (Raw data see other files).

#### 3.2 Results

The detection concentrations of phytohormones hormones in sample were calculated with the calibration cures. In summary, the content (ng/mL) of phytohormones was expressed as quantity of per volume (Raw and calculated data see other files). The contents of IAA in two samples (0.36 ng/mL vs 50.39 ng/mL) were the highest.

### 4 Conclusions and Discussions

Here we successfully performed ESI-HPLC-MS/MS analysis for plant hormones of 2 samples – BSF101E and FS101C - provided by customer. The concentration (ng/mL) of each item tested in each sample is listed in a summary table below (Table 3).

**Table 3. Summary of Contents**

| Component Name | BSF101E (ng/mL) | FS101C (ng/mL) |
|---|---|---|
| IAA | 0.36 | 50.39 |
| ABA | 0.039 | 5.37 |
| JA | 0.18 | 0.99 |
| Me-JA | 0.020 | 0.031 |
| GA1 | 0.068 | 0.025 |
| GA3 | 0.010 | 0.014 |
| GA4 | 0.0087 | 0.035 |
| GA7 | 0.00080 | 0.014 |

**– End of Report –**

Lifeasible

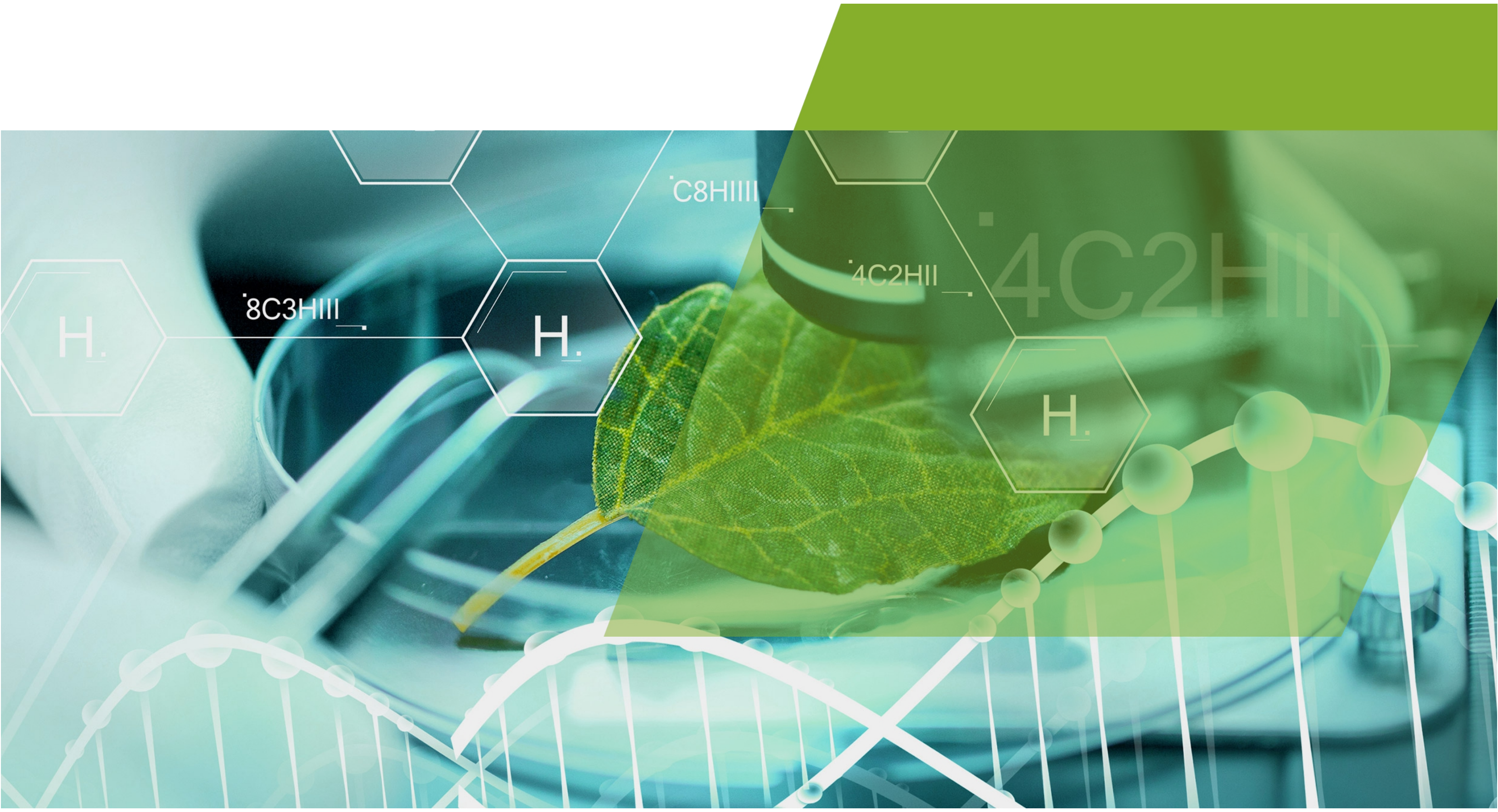

### CONTACT US

17 Ramsey Rd. Suite 209, SHirley, NY 11967

Web: https://www.lifeasible.com
