## Supplementary material for "A biochemical analysis of Black Soldier fly (*Hermetia illucens*) larval frass plant growth promoting activity": S3 HPLC GC-MS phytohormone analysis (from another version of this preprint: version 1, 2023-01-07)

| Sample name | Sample NO. | Diluted vol (mL) | Weighing vol (mL) |
|---|---|---|---|
| **BSF frass** | B1 | 0.200 | 5.000 |
| **Catering waste leachate -S1*** | B2 | 0.200 | 5.000 |
| **Catering waste leachate -S2*** | B2 | 0.200 | 5.000 |
| **Average S1, S2** | | | |

*S1 and S2 = duplicate samples analyzed by HPLC -GC-MS

| IAA | | MEJA | |
|---|---|---|---|
| Detection conc (ng/mL) | Content (ng/mL) | Detection conc (ng/mL) | Content (ng/mL) |
| 8.94 | **0.36** | 0.50 | **0.020** |
| 1279.01 | 51.16 | 0.77 | 0.031 |
| 1240.31 | 49.61 | 0.76 | 0.030 |
| | **50.39** | | **0.031** |

| ABA | | JA | |
|---|---|---|---|
| Detection conc (ng/mL) | Content (ng/mL) | Detection conc (ng/mL) | Content (ng/mL) |
| 0.97 | **0.039** | 4.44 | **0.178** |
| 135.19 | 5.41 | 24.92 | 0.997 |
| 133.14 | 5.33 | 24.44 | 0.978 |
| | **5.37** | | **0.99** |

| GA1 | | GA3 | |
|---|---|---|---|
| Detection conc (ng/mL) | Content (ng/mL) | Detection conc (ng/mL) | Content (ng/mL) |
| 0.17 | **0.0068** | 0.26 | **0.0104** |
| 0.62 | 0.025 | 0.34 | 0.014 |
| 0.65 | 0.026 | 0.34 | 0.014 |
| | **0.025** | | **0.014** |

| GA4 | | GA7 | |
|---|---|---|---|
| Detection conc (ng/mL) | Content (ng/mL) | Detection conc (ng/mL) | Content (ng/mL) |
| 0.22 | **0.0087** | 0.020 | **0.00080** |
| 0.87 | 0.035 | 0.34 | 0.0136 |
| 0.88 | 0.035 | 0.35 | 0.0141 |
| | **0.035** | | **0.014** |
